# Efficient CRISPR-Cas9-mediated genome editing of the cane toad (*Rhinella marina*)

**DOI:** 10.1101/2025.05.15.654396

**Authors:** Michael Clark, Alexander T. Funk, Alex Paporakis, Gregory P. Brown, Samuel J. Beach, Aidan Tay, Stephanie Deering, Caitlin Cooper, Mark Tizard, Chris J. Jolly, Georgia Ward-Fear, Anthony W. Waddle, Richard Shine, Maciej Maselko

## Abstract

Invasive species inflict major ecological, economic, social, and cultural harm worldwide, highlighting the urgent need for innovative and effective control strategies. Genome editing offers exciting possibilities for creating highly targeted control methods for invasive species. Here, we demonstrate CRISPR-Cas9 genome editing in the cane toad (*Rhinella marina*), one of Australia’s most notorious invasive species, by targeting the *tyrosinase* gene to produce albino phenotypes that provide clear visual markers for assessing editing efficiency. Microinjection of Cas9 protein and guide RNAs into one-cell zygotes resulted in 87.6% of mosaic larvae displaying nearly complete albinism, with 2.3% exhibiting complete albinism. For completely albino individuals, genomic analysis confirmed predominantly frameshift mutations or large deletions at the target site, with no wild-type alleles detected. Germline transmission rates reflected the extent of albinism in the mosaic adult, where we achieved maternal germline transmission rates of almost 100%. This technology, representing the first application of CRISPR-Cas9 in the Bufonidae family, opens possibilities for exploring both basic research questions and strategies for population control.

## Introduction

Invasive species are one of the leading drivers of global biodiversity loss and extinctions^1,2^ and impose a multi-trillion dollar economic burden^3^. Successful invaders often share traits such as fast life-history—characterised by rapid maturation, frequent reproduction, and high fecundity^4–6^—generalist ecology (i.e., broad diet and habitat use)^7,8^, behavioural flexibility,^9,10^ and broad environmental tolerance.^11,12^ These traits not only facilitate establishment and spread but also hinder efforts to control invasive species.^13,14^ Conventional approaches to invasive species management—such as trapping and manual removal—have often failed to deliver effective results, driving interest in more innovative control strategies.^15^ Advances in genetic technologies, particularly CRISPR-Cas9, have made it increasingly feasible to disrupt key gene pathways or introduce novel genetic material into invasive populations, offering a powerful new approach for population suppression.^16–18^ Most interest in using genetic modification for pest control has focused on transgenic approaches—introducing novel genetic material to modify pest animals in ways that reduce target populations.^19^ These methods range from modifications that sterilise males^20^ or make them reproductively toxic,^21^ to systems employing dominant lethal traits^22^ or self-propagating gene drives.^16^ Despite laboratory success, transgenic methods face significant regulatory hurdles globally, hindering their practical field application. The Australian context illustrates this: while discussions continue regarding potential field trials of transgenic mosquitoes, certain genetically modified organisms (GMOs) have already been approved, like commercially cultivated cotton, canola, and safflower.^23^ In contrast, gene knockouts that do not introduce novel DNA are not subject to the stringent regulations applied to GMOs. Therefore, we focus on this simpler and less controversial alternative—using CRISPR-Cas9 to knock out existing gene pathways without providing exogenous DNA as a repair template. This approach creates mutations that are indistinguishable from naturally occurring mutations. As a result such CRISPR-induced changes do not qualify as GMOs under Australian legislation.^24^

Cane toads (*Rhinella marina*), originally introduced to Queensland in 1935 to control cane beetles, have since become one of Australia’s most notorious invasive species^25^ and a prominent case study in invasion ecology and evolution.^26–28^ First introduced to sugar cane plantations near Gordonvale,^29,30^ the cane toad’s range now extends westwards almost to Broome, Western Australia and southwards to Grafton, New South Wales, encompassing over 1.2 million km^2^.^31^ Because Australia has no native bufonid (true toad) species, endemic lineages of frog-eating predators have no evolutionary history of exposure to the distinctive bufotoxin chemical defences of cane toads.^25^ As a result, toxic invasive toads have caused dramatic population declines in vulnerable native predators, including northern quolls (*Dasyurus hallucatus*;^32^ freshwater crocodiles (*Crocodylus johnstoni*;^33^, snakes (e.g., *Acanthophis* spp., *Pseudechis* spp.;^34,35^ and lizards (*Varanus* spp. and *Tiliqua scincoides intermedia*;^36,37^. The high fecundity of cane toads (to > 30,000 eggs per clutch^29^) renders direct culling an ineffective control over most of the species’ invaded range.^38,39^

There are two promising approaches for using gene knockouts via CRISPR-Cas9 to manage cane toads. The first targets the toxin-producing pathway; a CRISPR-modified toad that produces a less powerful toxin would pose less risk to vulnerable native predators, while still being distasteful enough to induce conditioned taste aversion to discourage the predator from eating more toads.^32,40^ The second approach, recently suggested by,^41^ exploits the cannibalistic behaviour of cane toad tadpoles. In Australia, cane toad tadpoles are highly cannibalistic—readily consuming toad eggs, but not those of native frogs.^42,43^ This selective behaviour has inspired a novel control strategy: using toad tadpoles to eliminate toad eggs.^41^ If metamorphosis could be blocked through CRISPR-Cas9 gene knockouts, a cohort of non-metamorphosing, egg-consuming tadpoles could persist in waterbodies and suppress toad recruitment.

The first step toward translating these ideas into on-the-ground management is to demonstrate that we can use CRISPR-Cas9 to genetically modify phenotypic traits in this non-model amphibian species. The CRISPR-Cas9 system consists of two key components: the Cas9 endonuclease and a guide RNA (gRNA) that directs Cas9 to specific genomic targets ^44,45^. The Cas9-gRNA complex binds to target DNA sequences that are adjacent to a protospacer adjacent motif (PAM)—typically NGG for the commonly used *Streptococcus pyogenes* Cas9—providing further sequence specificity.^46^ When bound, Cas9 creates a double-stranded break that is repaired by the cell’s native repair mechanisms, typically resulting in small insertions or deletions (indels) at the cut site.^47,48^ If these indels occur in coding regions and shift the reading frame, they can alter downstream amino acid sequences and introduce premature stop codons, disrupting protein function.^49^ This precise targeting ability makes CRISPR-Cas9 an ideal tool for gene knockout studies.^50,51^

CRISPR-Cas9 has been successfully deployed to disrupt genes in various amphibian species, including the frogs *Xenopus laevis*^52^ and *Xenopus tropicalis*,^53^ the salamanders *Pleurodeles waltl*^54^ and *Ambystoma mexicanum*^55^. Despite successful applications across several amphibian families, CRISPR-Cas9 genome editing has not been implemented in any species within the Bufonidae family. Our success would extend reverse-genetics approaches to true toads, enabling the study of gene function through targeted disruption. In other amphibian models, CRISPR-Cas9-mediated disruption for some genes has approached 100% efficiency ^53,55^, allowing researchers to study effects in the first generation without waiting for germline transmission to establish homozygous strains. For example, knockout studies identified the genes *catalase* and *fetub* as being important for limb regeneration in axolotls.^56^ Successes such as this across phylogenetically diverse amphibians helped inform our approach in adapting this technology to the cane toad.

Building on these advances in amphibian genome editing, we hope to establish CRISPR-Cas9 as a tool for both basic research on cane toad ecology and evolution and potential population control applications by demonstrating that we can efficiently edit the toad’s genome. We targeted the *tyrosinase* gene, which encodes an enzyme essential for melanin production in amphibians.^57^ This gene was selected because mutations in *tyrosinase* cause albinism, a recessive trait that is overt and provides rapid visual confirmation of successful bi-allelic editing across all developmental stages.^58^ We determined CRISPR-Cas9 disruption efficiency in larvae, assessed phenotypic penetrance across development, and evaluated germline transmission rates. These parameters are essential for future applications targeting genes involved in other biological processes such as metamorphosis and toxin production, whose disruption might facilitate control strategies for this invasive species.

## Materials and Methods

### Ethical statement

All experimental procedures involving animals were approved by the Macquarie University Animal Ethics Committee (approval number: 2023/015) and Institutional Biosafety Committee (approval number: 14368). Animals were euthanised with an overdose of pentabarbitone sodium (Lethabarb, Virbac, Milperra, Australia) in accordance with approved protocols. All efforts were made to minimise animal suffering and to reduce the number of animals used.

### Cane toad collection and husbandry

Adult cane toads were collected from Leaning Tree Lagoon, Northern Territory before being taken to the Macquarie University Tropical Ecology Research Facility. Adult cane toads were also collected from Grafton, northern New South Wales and taken to Macquarie University’s Sydney campus.

Toads kept in captivity for longer than 24 hrs were housed in plastic enclosures (64.5 x 41.3 x 27.6 cm) with water and hide tubes at 28 ± 2°C with 12:12 hour light cycle and 70–80% humidity. Animals were fed crickets three times weekly with vitamin supplements (URS Ultimate Vitamins, sourced from Budget Pet Products, Brisbane, Australia) and mealworms once weekly.

Tadpoles were maintained at densities ≤1 tadpole/litre in 60 L containers with either bore water (NT) or filtered water (Sydney), with 30% water changes every third day. Tadpoles were fed *ad libitum* with a 3:1 mixture of algae wafers to fish flakes once daily. Developmental stages were recorded according to the Gosner staging system.^59^ Recently-metamorphosed toadlets were initially fed live termites, before being transitioned to a diet of fish pellets and dog biscuits.

### Experimental design and gRNA design

We selected *tyrosinase* as our initial target gene due to its well-documented phenotype in amphibian knockouts, which provides clear visual confirmation of successful editing.^57,58^ The sequence for the first exon of cane toad *tyrosinase* (GenBank accession KR012509.1) was obtained for gRNA design. Using E-CRISP software (http://www.e-crisp.org/E-CRISP/), we designed two gRNAs targeting the first exon of tyrosinase. The selected gRNA sequences were: gRNA1 (5’-GTCTGGGCTGATGGTGCGTT-3’) and gRNA2 (5’-GCGTTGATGATCGGGAAAAC-3’). Custom crRNA and universal tracrRNA, and Alt-R® S.p Cas9 Nuclease V3 protein were purchased from Integrated DNA Technologies (IDT, Coralville, IA, USA).

We selected tyrosinase as our initial target gene due to its well-documented phenotype in amphibian knockouts, which provides clear visual confirmation of successful editing.^57,58^

### Breeding procedures

Toads were induced to breed by injection of leuprorelin acetate (Lucrin, Abbot Australasia, Botany, New South Wales, Australia) diluted in Simplified Amphibian Ringers’ (SAR) solution (0.25 mg/mL). Females were injected with 0.75 mL and males were 0.25 mL. Breeding pairs were housed in tilted plastic containers with 10 cm water depth at one end and dry land at the other. Egg strings were collected <15 min after deposition and maintained in smaller plastic containers. To increase fertilisation, IVF was performed on eggs.

Following Wlizla et al.,^60^ a male was euthanised and the testes were extracted. A quarter of a testis was placed into a 1.5 mL microcentrifuge tube with 1 mL of 1x SAR and was crushed with a bent pipette tip to release sperm. A monolayer of eggs was laid onto a petri dish, and the 1 mL solution of crushed testis was added to the eggs and mixed. After 5 minutes, the dish was flooded with aged water and was let to sit for an additional 5 minutes.

To de-jelly the fertilised eggs, a fresh solution of 1% L-cysteine free base (CAS# 52–90-4, Sigma Aldrich) was prepared in aged water. Fertilised eggs were transferred to this solution and periodically swirled until eggs at the edge of the jelly coat began to separate from the larger strand. At this point, the eggs were transferred to a solution of aged water. The eggs were passed through at least 6 separate solutions of aged water to remove residual L-cysteine. De-jellied eggs were then suitable for microinjection. For all egg movements, wide-bore plastic pipettes were used to minimise mechanical damage to the eggs.

### CRISPR-Cas9 delivery

Microinjections were performed using a modified technique adapted from established amphibian CRISPR-Cas9 delivery protocols^53,55^, in which a micromanipulator (M3301R, World Precision Instruments, Sarasota, FL, USA) equipped with a Digital Multi-pressure Microinjector (Tritech Research, Cat. No. MINJ-D) was used. Quartz needles were pulled from quartz capillaries (1.0 mm O.D., 0.75 mm I.D.; Sutter Instrument Co.) using a Sutter P-2000 laser micropipette puller with the following parameter settings: Heat = 735, Filament = 4, Velocity = 50, Delay = 126, Pull = 120. Needle tips were bevelled at a 35° angle using a microgrinder (Narishige EG-45, Tokyo, Japan).

De-jellied eggs were immobilised in a custom injection dish with lanes formed in 3% agarose prepared in aged water. Needles were backfilled with the injection solution containing Cas9 protein (IDT, 200 ng/μL) and guide RNAs (15 ng/μL each) prepared in IDT duplex buffer. Each egg received two injections into the animal pole, with approximately 10 nL delivered per injection site (20 nL total per egg). Injections were performed within at most 60 min post-fertilisation to ensure delivery before first cleavage. Following injection, eggs were transferred to aged water and maintained at 28°C.

### Viability assessment

The impact of CRISPR-Cas9 genome editing on cane toad development was assessed by quantifying survival and morphological outcomes following Kroll et al.^50^. Embryonic viability was evaluated by tracking developmental progression from 1 day post-fertilisation (dpf) onward. Any embryos exhibiting mortality or abnormal development before 1 dpf were excluded from analysis, as these early failures typically represent fertilisation issues or mechanical trauma from the microinjection procedure rather than gene editing effects.

To account for background mortality not associated with CRISPR-Cas9 activity, control groups of embryos that were subjected to L-cysteine treatment but not injection. This background mortality rate was subtracted from the unviability rate in the injected groups to isolate effects specifically caused by the mutagenic ribonucleoprotein complex injections. For example, if 10% of the injected embryos died or were dysmorphic during the 1-4 dpf window, and 2% of the l-cysteine-treated control embryos died during the same period, the CRISPR-specific unviability rate was recorded as 8%. Tadpoles were monitored daily until 4 dpf, and a subset were then raised to sexual maturity for germline transmission assessment.

### Phenotype scoring

Phenotypic assessment of F0 animals was performed at multiple developmental stages. Following Kroll et al.^50^ at 4 days dpf, each animal was scored on a standardised 5-point scale based on eye pigmentation: score 1 represented complete absence of pigmentation; score 2 indicated minimal pigmentation with only one or two patches of pigment; score 3 showed approximately half-normal pigmentation; score 4 exhibited mostly normal pigmentation with small unpigmented areas; and score 5 displayed normal wild-type pigmentation. When animals had eyes with different pigmentation levels, the score of the more pigmented eye was recorded. All scoring was performed by an observer blinded to the experimental condition. Body pigmentation was also recorded for each animal, though the standardised eye scale provided a more discrete and easily scored phenotype compared to the complex mosaic patterns observed across the body. For later developmental stages (metamorphs and adults), animals were photographed from multiple angles to document the albino phenotype.

### Genomic DNA extraction

Genomic DNA was extracted using the High Pure PCR Template Preparation Kit (Roche Diagnostics) to assess CRISPR-induced mutations at the tyrosinase locus. For F0 analysis, whole 4 dpf tadpoles were used, while toe clips were collected from F1 heterozygotes. 4 dpf larvae were euthanised with MS-222 (tricaine methanesulfonate, 0.5 g/L, pH 7.5). Each sample was transferred to a 1.5 mL microcentrifuge tube and processed according to the manufacturer’s protocol.

Briefly, tissues were lysed in Tissue Lysis Buffer with Proteinase K (provided in the kit) and incubated at 55°C overnight. Lysates were combined with Binding Buffer and Isopropanol, then loaded onto the kit’s High Pure filter columns. DNA was bound to the silica membrane through centrifugation (13,000 × g for 1 minute), followed by two washes with Wash Buffer. Purified DNA was eluted in molecular grade water and quantified using a NanoDrop spectrophotometer (Thermo Fisher Scientific).

### Sanger sequencing

To analyse editing at the *tyrosinase* locus, PCR amplicons spanning both target sites were generated using forward primer (5’-CCTGGAGGAGTAATAGTGAAGCC-3’) and reverse primer (5’-TCCTTGCGGAGAAGCTTCTG-3’). PCR was performed using Q5 High-Fidelity DNA Polymerase (NEB). PCR products were purified using the QIAquick PCR Purification Kit (QIAGEN, Hilden, Germany; Cat. No. 28104) according to the manufacturer’s instructions and subjected to Sanger sequencing at the Australian Genome Research Facility (AGRF). For each sample, bidirectional Sanger sequencing was performed using PCR amplicons spanning both target sites. The forward primer was used to analyse editing at the gRNA1 target site, while the reverse primer was used to analyse editing at the gRNA2 target site. Sequence analysis was performed using the DECODR (v3.0) online tool (https://decodr.org/). For each sample, a knockout (KO) score was calculated, representing the percentage of frameshift mutations (indels not divisible by three) or large deletions (≥21 bp) that would be expected to disrupt protein function. Additionally, the frequency and specific characteristics (insertion/deletion size and type) of each mutation were quantified and catalogued for each sample. All analysed samples, unless indicated, had DECODR r² values greater than 0.6, indicating reliable trace decomposition.

### Measurement of germ line transmission

To assess germline transmission of CRISPR-induced mutations, we raised edited F0 animals to sexual maturity. Two crosses were established between F0 animals showing varying degrees of albinism. For Cross 1, a completely albino female (**Figure 2B**) was paired with a mosaic male (**Supplementary Figure 2**), while Cross 2 paired a completely albino female (**Figure 4B**) with a mosaic male (**Figure 4A**).

To induce breeding, adults were injected with leuprorelin acetate as described above. Egg strings were collected and maintained in separate containers. Phenotypic scoring of eggs and resulting tadpoles was performed to determine maternal and paternal transmission rates of the albino trait. Eggs were scored as albino when they lacked all visible pigmentation, and wild-type when normal pigmentation was present. For each cross, a subset of eggs were raised to assess larval phenotypes and confirm germline transmission.

## Results

### Survival and phenotypic characterisation of F0 albino and mosaic tadpoles

CRISPR-Cas9 modification of the *tyrosinase* gene resulted in high-efficiency phenotypic changes in cane toad embryos (**Figure 1A)**. Across six clutches, 87.6% of larvae (304/347) displayed nearly complete albinism (score 2), with 8 larvae (2.3%) exhibiting complete albinism (score 1) and only 19 (5.5%) retaining wild-type pigmentation (score 5). The remaining 16 larvae (4.6%) showed intermediate pigmentation patterns (score 3–4). In contrast, all uninjected control larvae (n = 734) displayed wild-type pigmentation (score 5). Post-injection survival rates varied between clutches, ranging from 60-97% (**Supplementary Figure 1**).

**Figure 1:**
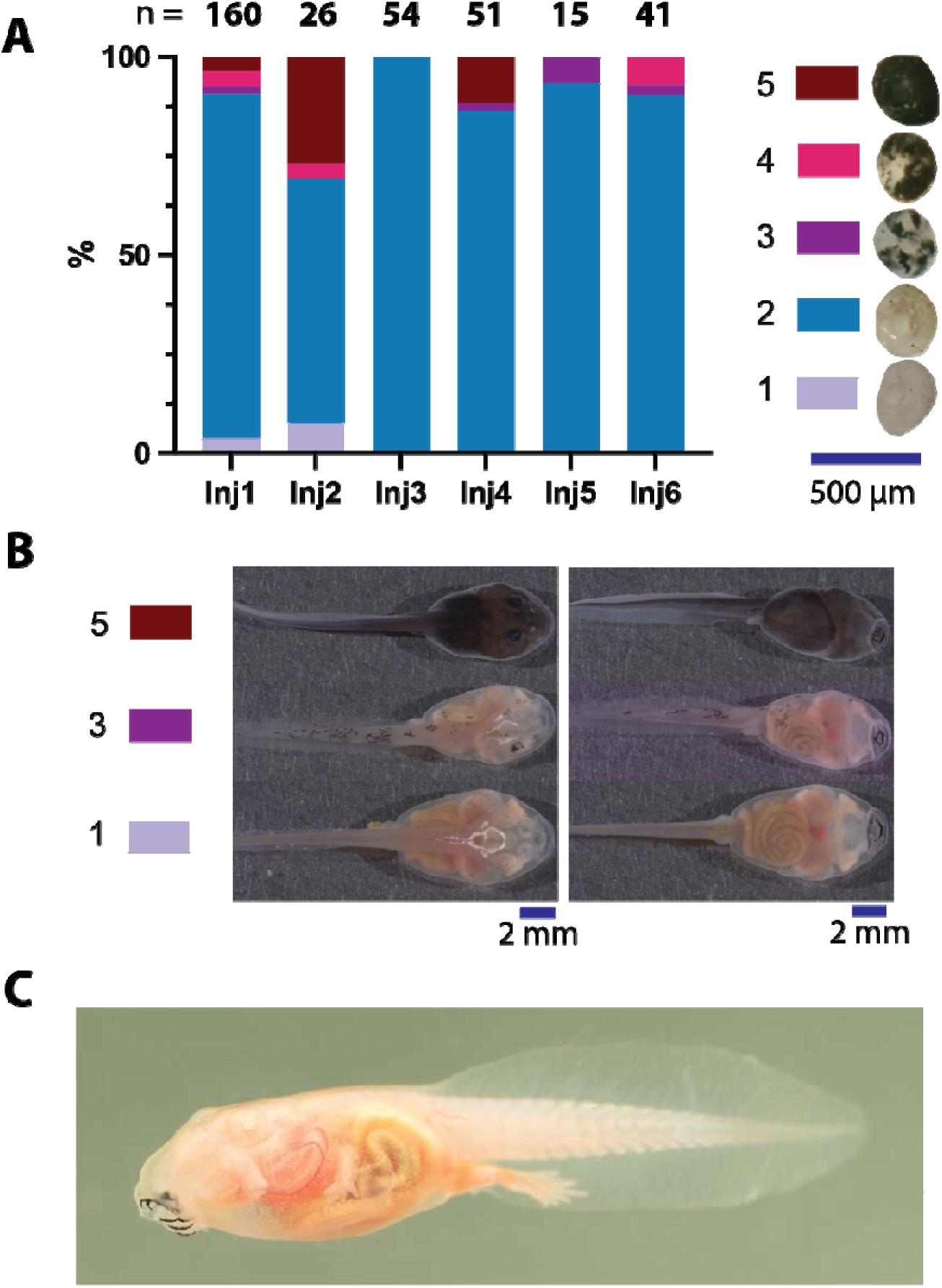
F0 albino tadpole phenotyping data. A) Six injection sessions were conducted, and eye phenotypes were scored based on the level of pigmentation, with representative images shown on the right. Eyes at 4 days post-fertilisation (dpf) range from score 1 (complete absence of pigment, shown in light purple) to score 5 (fully pigmented wild-type, shown in maroon). The colour coding of the bars in the graph corresponds to the same pigmentation level categories shown in the images. Sample sizes (n) for each injection group are indicated at the top of each column. B) Representative images of tadpoles at Gosner stages 30-33 with both dorsal and ventral views, exhibiting different pigmentation phenotypes. Top row shows score 5 tadpoles, middle row shows score 3 tadpoles and bottom row shows score 1 tadpoles. C) An F0 score 1 tadpole at Gosner stage 39, allowing observation of its internal organs, including the gut, heart and developing lungs. Tadpole photograph by Etienne Littlefair, with permission.

Body pigmentation largely mirrored eye pigmentation, with albino individuals also lacking melanin in their skin, allowing clear visualisation of internal organs (**Figure 1B**, bottom row and **Figure 1C**). Partially edited animals displayed mosaic patterns, with patches of pigmented and unpigmented tissue distributed across the body (**Figure 1B**, middle row). Although these body pigmentation patterns were recorded (see **Supplementary file 1**), the standardised 5-point eye pigmentation scale provided a more discrete and easily scored phenotype.

### Persistence of albino phenotypes through metamorphosis and into adulthood

The pigmentation phenotypes observed at the tadpole stage persisted into the metamorph stage (**Figure 2A**) and into adulthood (**Figure 2B**). Animals that exhibited fully penetrant albino phenotypes as tadpoles maintained their lack of pigmentation as metamorphs and adults, whereas individuals with mosaic patterns retained their patchy pigmentation distribution. This persistence of phenotypes across developmental stages demonstrates stable disruption of the *tyrosinase* gene in affected tissues throughout the cane toads’ life cycle.

**Figure 2:**
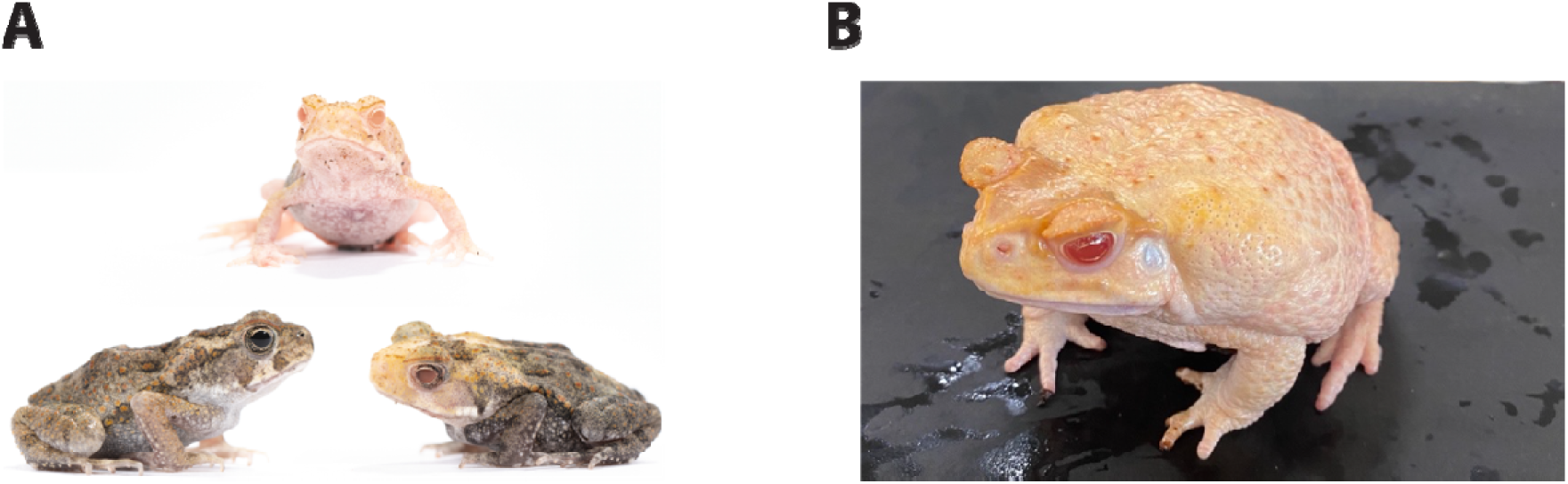
Penetrance of the *tyrosinase* mutation in metamorphosed toads. A) Three F0 metamorphs displaying different pigmentation phenotypes. Top: an F0 albino metamorph with no visible dark pigmentation; Bottom: two metamorphs showing wild-type (left) and mosaic pigmentation patterns (right) with patches of pigmented and unpigmented tissue. B) An F0 adult female toad with no detectable dark pigmentation. Metamorph photographs by Etienne Littlefair, with permission.

### Molecular characterisation of CRISPR-induced *tyrosinase* mutations

To show that the observed phenotypes resulted from CRISPR-mediated mutations in the *tyrosinase* gene, we performed genomic DNA analysis on F0 albino and partially pigmented larvae at 4 dpf. PCR amplicons spanning both gRNA target sites were generated and subjected to Sanger sequencing (**Figure 3A**). High-confidence sequencing data (DECODR r^2^ > 0.6) was obtained for 4 albino samples and 4 partially albino samples t both gRNA1 and gRNA2.

**Figure 3:**
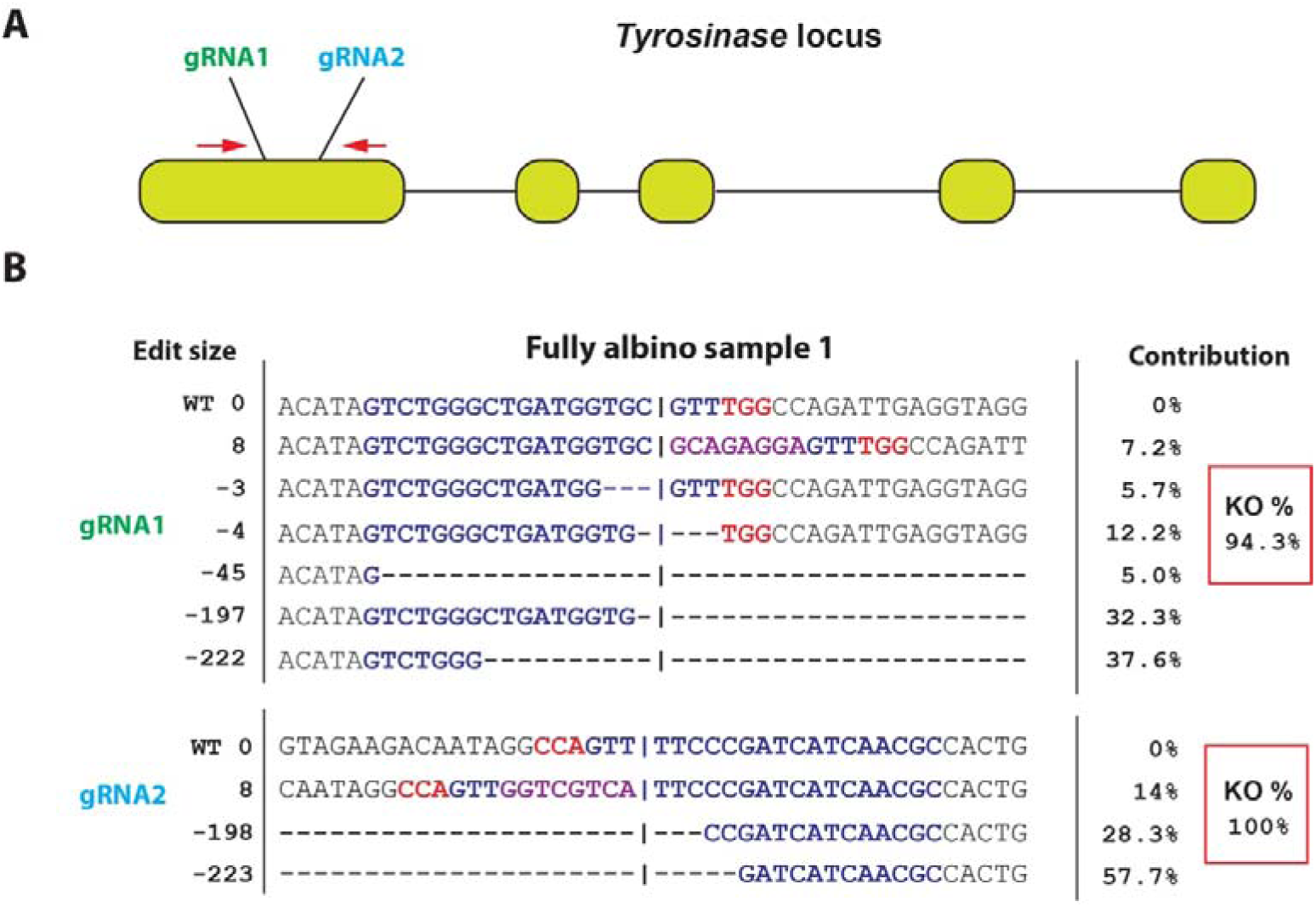
Molecular characterisation of CRISPR/Cas9-induced mutations of the *tyrosinase* gene. A) Schematic representation of the *tyrosinase* locus showing the locations of gRNA1 (green) and gRNA2 (light blue) target sites along with the PCR primers (red) used for mutational analysis. B) Sequence analysis at both gRNA sites for a fully albino F0 tadpole, as determined using the DECODRv3.0 analysis tool. For each target site, the gRNA sequence is indicated by dark blue text, with the PAM sequence in red. The vertical line in the text marks the Cas9 cut site. Below each reference sequence, individual mutation types are shown with their respective contributions (%). Deletions are represented by dashes while insertions are shown in purple text. The knockout (KO) scores (red boxes) indicate the percentage of mutations causing frameshift or large deletions (≥21 bp) at each site. At gRNA1, 94.3% of detected sequences were knockout mutations, while at gRNA2, 100% of sequences were knockout mutations. The absence of wild-type sequences at both target sites confirms efficient bi-allelic disruption of the *tyrosinase* gene, corresponding with the complete albino phenotype.

In completely albino larvae, sequencing revealed complete sequence disruption beginning at the Cas9 cut sites (**Figure 3B**). At the gRNA1 target site, three of four samples with high-confidence sequencing data (r^2^ > 0.6) exhibited 100% KO scores, with the remaining sample showing a 94.3% score (**Supplementary table 1**). The top panel of **figure 3B** illustrates the diversity of mutations detected at the gRNA1 site in the 94.3% KO sample, showing various deletions and insertions. In this sample, most mutations (94.3%) were frameshift indels or large deletions that would disrupt protein function, with only 5.7% showing a 3 bp in-frame deletion. The mutations across all samples primarily consisted of small indels (1-17 bp) and larger deletions (up to 222 bp), with no wild-type alleles detected in any of the completely albino samples.

Sequencing analysis at the gRNA2 target site also showed complete disruption of wild-type sequences in all completely albino larvae. Among the four samples with high-confidence data (r^2^ > 0.6), KO scores ranged from 78.8% to 100%. Despite the variable KO scores, no wild-type sequences were detected at gRNA2 in any of these completely albino individuals, suggesting highly efficient Cas9 cutting, but with different repair outcomes. The most common mutations observed were small indels (1-13 bp) and larger deletions (up to 223 bp) that spanned both gRNA sites.

Partially albino individuals displayed notably different mutation profiles. KO scores ranged from 3.5% to 53.6% at gRNA1 and from 8.8% to 58.5% at gRNA2, with wild-type alleles comprising 40.1–96.5% and 41.5-73.2% of sequences, respectively. The mutations observed in these mosaic animals were predominantly small indels up to 22 bp, with no detections of large deletions spanning both gRNA sites. The strong correlation between the molecular data and phenotypic observations demonstrates that the albino phenotypes resulted from bi-allelic disruption of the *tyrosinase* gene. Complete albinism occurred when all alleles were frameshift mutations or large deletions at one of the gRNA sites, whereas partial albinism resulted from mosaic editing where significant proportions of wild-type alleles remained.

### Germline transmission of *tyrosinase* mutations

Maternal transmission rates reached or approached 100% in both crosses, with Cross 1 producing 99.93% albino eggs (1545/1546) and Cross 2 yielding 100% albino eggs (498/498) (Table 1, Cross 1 eggs: **Figure 4C**, Cross 2 eggs: **Figure 4D**). This maternal transmission rate reflects the near-complete albinism observed in both F0 mothers (Cross 1 mother: **Figure 2B**, Cross 2 mother: **Figure 4B**).

**Figure 4:**
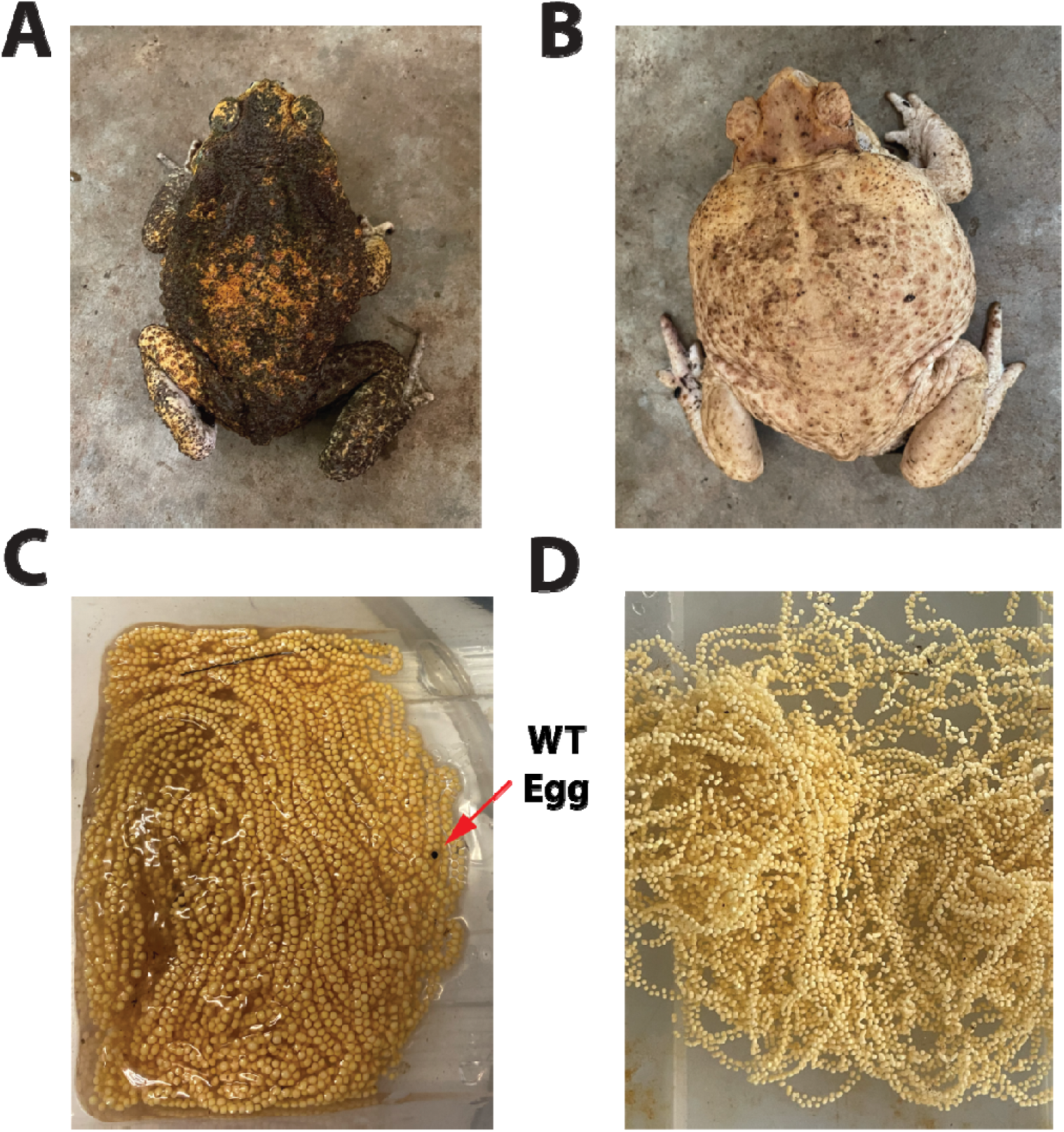
Germline transmission of CRISPR-induced *tyrosinase* mutations. A) Adult F0 male from Cross 2; it is partially albino with a mosaic pigmentation pattern. B) Adult F0 female toad from Cross 2 with complete albinism, displaying no visible dark pigmentation. C) Clutch of eggs from Cross 1 showing nearly 100% albino phenotype (1545/1546 eggs– only a subset was counted), demonstrating highly efficient maternal transmission of the *tyrosinase* mutation. The single, darkly pigmented, WT egg is indicated with a red arrow. D) Clutch of eggs from Cross 2 showing 100% albino phenotype (498/498 eggs – only a subset was counted). The high efficiency of maternal germline transmission observed in both crosses corresponds with the complete or nearly complete albino phenotype of the F0 mothers.

**Table 1:** The germline transmission of albinism in F1 offspring. Albinism was recorded for two crosses between a total of four F0 albino parents.

| Cross | Albino<br>eggs | WT<sup>a</sup><br>eggs | % Maternal<br>transmission | Albino<br>larvae | WT larvae | % Paternal<br>transmission |
| --- | --- | --- | --- | --- | --- | --- |
| 1 | 1545 | 1 | 99.93 | 1 | 2 | 33 |
| 2 | 498 | 0 | 100 | 115 | 383 | 23.1 |

<sup>a</sup>WT = wild-type

Paternal transmission rates were substantially lower. Only 33% (1/3) and 23.1% (115/498) of fertilised eggs inherited the albino phenotype from the father in Cross 1 and Cross 2, respectively. This lower transmission rate corresponds with the more mosaic phenotype observed in the F0 fathers (Cross 2 father: **Figure 4A**, Cross 1 father: **Supplementary Figure 2**), indicating less extensive editing of the germline. The very low number of larvae for Cross 1 may have been due to low egg viability or low rate of fertilisation. All phenotypically albino F1 offspring exhibited compound heterozygosity at the gRNA2 target site, with complete absence of wild-type sequences (Figure 5). Six of 8 samples carried an in-frame 3-bp deletion that removed an asparagine residue.

**Figure 5:**
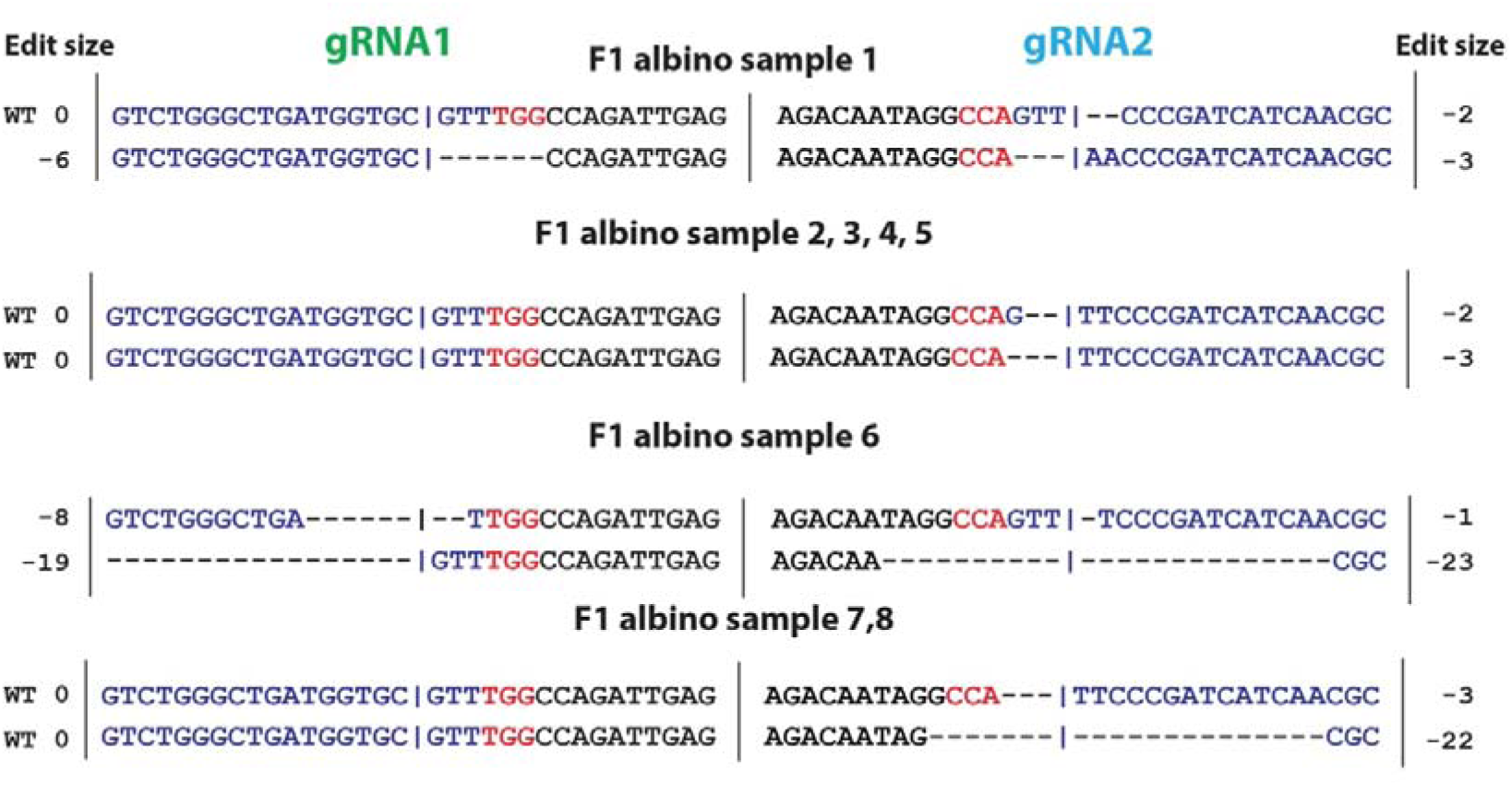
Molecular analysis of F1 albino offspring showing transmitted CRISPR-induced mutations in the tyrosinase gene. The figure shows sequence analysis of eight F1 albino samples at both gRNA target sites. For each target site, the gRNA sequence is indicated by dark blue text, with the PAM sequence in red. The vertical line in the text marks the Cas9 cut site. Deletions are represented by dashes. At the gRNA1 site, samples exhibited various editing patterns including wild-type sequence retention and deletions ranging from −6 to −19 bp. At the gRNA2 site, all samples showed bi-allelic disruption with no wild-type sequences detected. Notably, samples 1-5, 7, and 8 carried a minimal 3-bp deletion at the gRNA2 site that removes a single asparagine residue. These results demonstrate that even small mutations can be stably transmitted through the germline while producing complete phenotypic effects.

## Discussion

Our results demonstrate the first successful application of CRISPR-Cas9 genome editing in the Bufonidae family, achieving high editing efficiency in cane toads. Following editing at the *tyrosinase* gene, 87.6% of F0 larvae displayed nearly complete albinism as measured by eye pigmentation, while 2.3% exhibited complete albinism. This high success rate in a non-model amphibian species was validated by genomic sequencing, which showed predominantly frameshift mutations (1–17 bp indels) and larger deletions (up to 223 bp) at the target sites, with no wild-type alleles detected in completely albino larvae. We found that even subtle alterations to the amino acid sequence affected enzyme function—as demonstrated in the F1 offspring, where the in-frame deletion of a single asparagine residue was enough to disrupt tyrosinase activity and produce albinism. Importantly, we observed successful germline transmission, with maternal transmission rates approaching 100%. The albino phenotype persisted across all developmental stages from tadpole to adult, confirming the stability of the gene knockout throughout development.

The editing efficiency recorded in this study is comparable to that recorded in other CRISPR-Cas9 applications in amphibian species, including *Xenopus* and axolotls, where editing rates also approach 100% for some target genes.^52,53,55^ The ability to generate high-efficiency knockouts in F0 animals eliminates the need to wait for subsequent generations to observe phenotypic effects.^50,61^ This is particularly valuable for cane toads, which have a relatively long generation time (∼9 months) relative to model organisms. Rather than requiring a stable knockout line, F0 knockouts that approach 100% efficiency can be studied immediately, providing rapid insights into gene function.

Beyond these technical advantages, our approach can provide direct causal evidence for the genetic underpinnings of variation in phenotypic traits within a population. Extensive research in evolutionary biology is based on the concept of genetic variation driving phenotypic variation that in turn affects an organism’s viability (evolutionary fitness).^62^ However, almost all of this research relies upon phenotypic variation whose genetic basis is unclear.^27,31,63^ CRISPR-Cas9 technology overcomes this limitation by enabling targeted genetic modifications to directly measure resulting phenotypic effects and their impact on fitness. For example, the targeted knockout of Heat Shock Transcription Factor 1 in corals established its role in thermal tolerance.^64^ The Australian cane toad invasion provides an ideal system for deploying this approach, given the species’ well-documented rapid adaptation across diverse environments and the identification of candidate genes potentially under selection, such as Heat Shock Proteins.^26,65^ By causally inducing phenotypic change through gene knockouts, we can obtain more robust insights into the ways in which mutation affects microevolutionary fitness, and hence gene-frequency changes through time and space.^66,67^

The successful application of CRISPR-Cas9 in cane toads also provides new possibilities for invasive species management. One particularly promising direction is the potential to create tadpoles that remain in the larval stage indefinitely by targeting genes essential for metamorphosis. As proposed by Shine & Baeckens,^41^ non-metamorphosing tadpoles could be added to breeding sites, where they would cannibalise newly laid toad eggs, chemically suppress hatchlings and compete with post-hatchling larvae. These non-metamorphosing tadpoles would pose minimal threat to native fauna while effectively suppressing toad reproduction.^42,43^ Our demonstration that CRISPR-Cas9 can efficiently disrupt gene function in cane toads is an important step toward developing this novel control strategy. This technology also furthers the possibility of targeting specific genes to create toads with reduced toxicity to diminish their impact on native predators.^15,40^ Importantly, in many jurisdictions, gene knockouts created through CRISPR-Cas9 without the introduction of foreign genetic material are not classified as GMOs, which streamlines approval processes for field testing^24^ and allows for the possibility of releasing knockouts *in situ* for pest control.

Here, we have established a technique for the genetic modification of cane toads that will enhance our ability to study the basic biology of this invasive species while also facilitating the development, testing, and potentially implementation of new population control strategies. The high editing efficiency and successful germline transmission observed in our study demonstrates that CRISPR-Cas9 can be deployed effectively and efficiently in this non-model amphibian species. This work sits at the intersection of molecular biology and conservation genetics and offers new tools to address one of Australia’s most significant conservation challenges.

## Acknowledgements

We thank Stewart Macdonald for his assistance with toad collection. We thank the Macquarie University Animal Services (MARS) staff for expert animal care, particularly Jason Martin-Powell, Ronny Eidels Shimonny, Natalie Casha, Bridget Butler, Natasha Suhochev, Alora Cantwell, Anna Lukan, Cheryl Song, Ayesha Earey, Tonje Skar, Taylah Portelli, Emelie Toovey, Chris Wilson and Josh Tsui. Special thanks to Jamie Harding for help with toad husbandry at Macquarie University’s Tropical Ecology Research Facility. We are grateful to Jason Martin-Powell and Melanie Elphick for their help in establishing the Macquarie University toad facility. We are also grateful to Etienne Littlefair for providing the photographs of the albino tadpole and metamorphs. This research was supported by the Minderoo Foundation (Grant: CLB-2372).

## Authorship Confirmation Statement

All authors have read and agreed to the published version of the manuscript. All authors have made substantial contributions to the work reported and agree to be accountable for all aspects of the research. The manuscript has been approved by all named authors for publication.

## Supplementary information

**Supplementary Figure 1:**
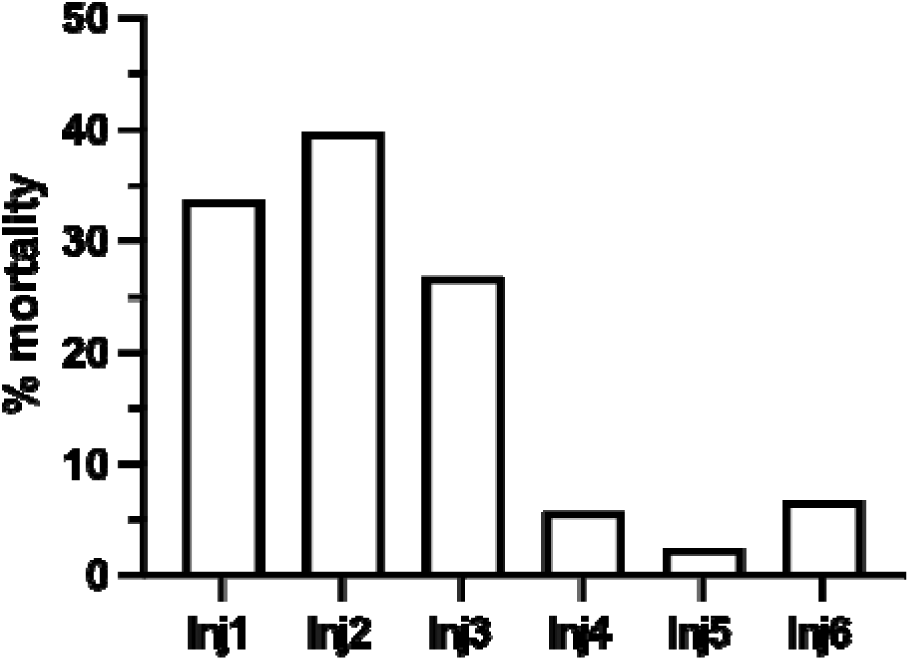
Embryo viability across six CRISPR injection sessions. The graph shows the percentage of unviable embryos for each tyrosinase (Tyr) injection session. Sample sizes (n) for each injection group are indicated at the top of the figure.

**Supplementary Figure 2:**
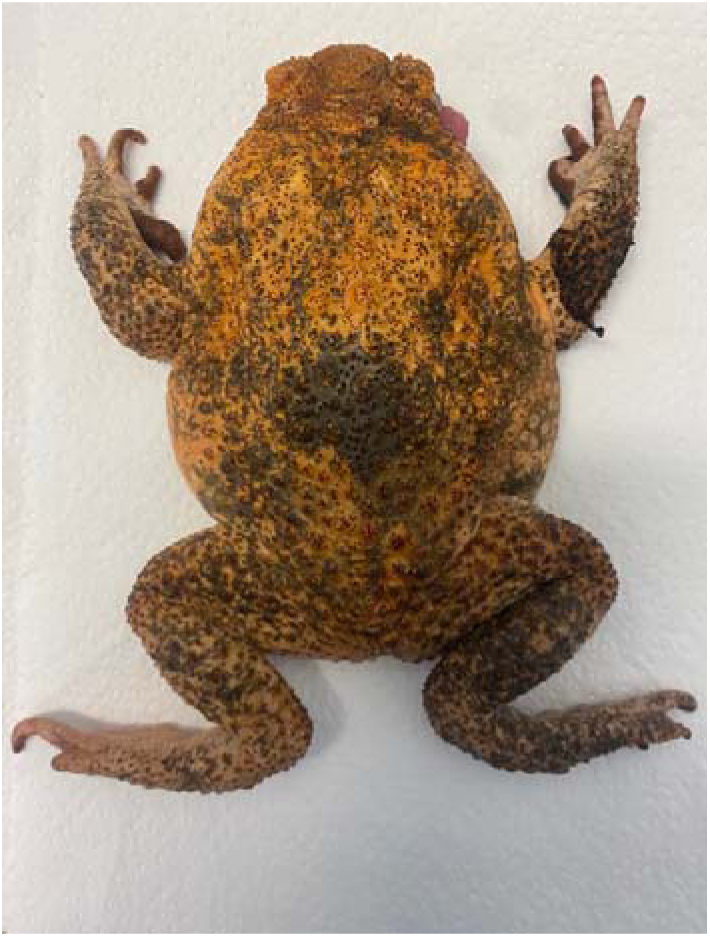
Cross 1 paternal toad. Dorsal view of the partially albino F0 male used in Cross 1, displaying a mosaic pigmentation pattern. This paternal mosaic phenotype correlates with the lower transmission rate (33%) of the albino trait observed in offspring.

**Supplementary table 1:** Summary of editing at the *tyrosinase* locus in albino and partially albino tadpoles. The table shows sequence analysis results from 5 fully albino samples and 3 partially albino samples at both gRNA target sites. For each sample, knockout (KO) percentages, edit sizes, and relative contributions are presented. For each target site and sample, the KO score represents the percentage of frameshift mutations or large deletions (≥21 bp). Edit sizes are shown as deletions (negative numbers) or insertions (positive numbers with + prefix), with ‘0 (WT)’ indicating wild-type sequence. Values in red text indicate data derived from lower confidence sequencing results, with their r² value detailed below the KO value.

| Sample | gRNA1 |  |  | gRNA2 |  |  |
| --- | --- | --- | --- | --- | --- | --- |
|  | KO % | Edit size | Contribution % | KO % | Edit size | Contribution % |
| Fully albino 1 | 94.3 | -222 | 37.6 | 100 | -223 | 57.7 |
|  |  | -197 | 32.3 |  | -198 | 28.3 |
|  |  | -4 | 12.2 |  | +8 | 14 |
|  |  | +8 | 7.2 |  |  |  |
|  |  | -3 | 5.7 |  |  |  |
|  |  | -45 | 5 |  |  |  |
| Fully albino 2 | 100 | -4 | 51.2 | 78.8 | -13 | 40.9 |
|  |  | +4 | 28.2 |  | -9 | 11.1 |
|  |  | -206 | 11.2 |  | -12 | 9.2 |
|  |  | -17 | 9.4 |  | -10 | 8.7 |
|  |  |  |  |  | -4 | 7.7 |
|  |  |  |  |  | +2 | 6.5 |
|  |  |  |  |  | -2 | 5.7 |
|  |  |  |  |  | -42 | 5.4 |
|  |  |  |  |  | -3 | 4.9 |
| Fully albino 3 | 100 | -197 | 50.6 | 78.6<br>($r^2=0.59$) | -197 | 38.4 |
|  |  | -17 | 44.7 |  | -9 | 21.4 |
|  |  | -7 | 4.7 |  | -8 | 16.6 |
|  |  |  |  |  | -196 | 10.4 |
|  |  |  |  |  | -185 | 8 |
|  |  |  |  |  | +4 | 5.1 |
| Fully albino 4 | 100 | -17 | 60.2 | NA<sup>a</sup> |  |  |
|  |  | -7 | 14.4 |  |  |  |
|  |  | -195 | 14.4 |  |  |  |
|  |  | +7 | 10.9 |  |  |  |
| Fully albino 5 | 100 | -208 | 75 | 89.8 | -208, 1 | 63.7 |
| | ($r^2=0.59$) | -17 | 24.9 | | -9 | 10.2 |
|  |  |  |  |  | -3, 1 | 9.2 |
|  |  |  |  |  | -9, 1 | 6 |
|  |  |  |  |  | -4, 3 | 6 |
|  |  |  |  |  | -39, 1 | 5 |
| **Fully albino 6** | 81.1 | -208 | 59.3 | 81.7 | -210 | 61.5 |
| | ($r^2=0.51$) | -3 | 18.9 | | -12 | 18.3 |
|  |  | -40 | 12.8 |  | -2 | 16.8 |
|  |  | -87 | 9.1 |  | -33 | 3.4 |
|  |  |  |  |  |  | -33 |
| **Partial albino 1** | 53.6 | 0 (WT<sup>b</sup>) | 40.1 | 58.5 | 0 (WT) | 41.5 |
|  |  | -4 | 34.2 |  | -2 | 40.4 |
|  |  | -17 | 9.2 |  | -21 | 9.6 |
|  |  | -7 | 6.6 |  | -8 | 8.5 |
|  |  | -6 | 4.2 |  |  |  |
|  |  | -9 | 2 |  |  |  |
|  |  | +1 | 2 |  |  |  |
|  |  | -8 | 1.6 |  |  |  |
| **Partial albino 2** | 3.5 | 0 (WT) | 96.5 | 8.8 | 0 (WT) | 73.2 |
|  |  | -4 | 3.5 |  | -3 | 18 |
|  |  |  |  |  | -22 | 4.6 |
|  |  |  |  |  | -2 | 4.2 |
| **Partial albino 3** | 32.2 | 0 (WT) | 67.8 | 20.9 | 0 (WT) | 79.1 |
|  |  | -4 | 18.2 |  | -2 | 14 |
|  |  | 7 | 14 |  | -8 | 6.9 |
| **Partial albino 4** | 18.6 | 0 (WT) | 81.4 | 25.4 | 0 (WT) | 64.5 |
|  |  | -4 | 18.6 |  | -2 | 24.1 |
|  |  |  |  |  | -9 | 10.1 |
|  |  |  |  |  | -8 | 1.4 |

<sup>a</sup>NA indicates that no sequence data was obtained for that sample at the respective gRNA site.
<sup>b</sup>WT= wild-type

## References

1. Bellard C, Cassey P, Blackburn TM. Alien species as a driver of recent extinctions. Biology Letters 2016;12(2):20150623; doi: 10.1098/rsbl.2015.0623.

2. Clavero M, García-Berthou E. Invasive species are a leading cause of animal extinctions. Trends in Ecology & Evolution 2005;20(3):110; doi: 10.1016/j.tree.2005.01.003.

3. Diagne C, Leroy B, Vaissière A-C, et al. High and rising economic costs of biological invasions worldwide. Nature 2021;592(7855):571–576; doi: 10.1038/s41586-021-03405-6.

4. Allen WL, Street SE, Capellini I. Fast life history traits promote invasion success in amphibians and reptiles. Ecol Lett 2017;20(2):222–230; doi: 10.1111/ele.12728.

5. Capellini I, Baker J, Allen WL, et al. The role of life history traits in mammalian invasion success. Ecology Letters 2015;18(10):1099–1107; doi: 10.1111/ele.12493.

6. Sol D, Maspons J, Vall-llosera M, et al. Unraveling the life history of successful invaders. Science 2012;337(6094):580–583; doi: 10.1126/science.1221523.

7. Gallagher RV, Randall RP, Leishman MR. Trait differences between naturalized and invasive plant species independent of residence time and phylogeny. Conservation Biology 2015;29(2):360–369; doi: 10.1111/cobi.12399.

8. Marchetti MP, Moyle PB, Levine R. Invasive species profiling? Exploring the characteristics of non-native fishes across invasion stages in California. Freshwater Biology 2004;49(5):646–661; doi: 10.1111/j.1365-2427.2004.01202.x.

9. Chapple DG, Simmonds SM, Wong BBM. Can behavioral and personality traits influence the success of unintentional species introductions? Trends in Ecology & Evolution 2012;27(1):57–64; doi: 10.1016/j.tree.2011.09.010.

10. Sol D, Timmermans S, Lefebvre L. Behavioural flexibility and invasion success in birds. Animal Behaviour 2002;63(3):495–502; doi: 10.1006/anbe.2001.1953.

11. Kolar CS, Lodge DM. Progress in invasion biology: predicting invaders. Trends in Ecology & Evolution 2001;16(4):199–204; doi: 10.1016/S0169-5347(01)02101-2.

12. Thuiller W, Richardson DM, Pyšek P, et al. Niche-based modelling as a tool for predicting the risk of alien plant invasions at a global scale. Global Change Biology 2005;11(12):2234–2250; doi: 10.1111/j.1365-2486.2005.001018.x.

13. Mack RN, Simberloff D, Mark Lonsdale W, et al. Biotic invasions: causes, epidemiology, global consequences, and control. Ecological Applications 2000;10(3):689–710; doi: 10.1890/1051-0761(2000)010[0689:BICEGC]2.0.CO;2.

14. Simberloff D, Martin J-L, Genovesi P, et al. Impacts of biological invasions: what’s what and the way forward. Trends Ecol Evol 2013;28(1):58–66; doi: 10.1016/j.tree.2012.07.013.

15. Tingley R, Ward-Fear G, Schwarzkopf L, et al. New weapons in the toad toolkit: a review of methods to control and mitigate the biodiversity impacts of invasive cane toads (*Rhinella marina*). The Quarterly Review of Biology 2017;92(2):123–149; doi: 10.1086/692167.

16. Champer J, Buchman A, Akbari OS. Cheating evolution: engineering gene drives to manipulate the fate of wild populations. Nat Rev Genet 2016;17(3):146–159; doi: 10.1038/nrg.2015.34.

17. Champer J, Lee E, Yang E, et al. A toxin-antidote CRISPR gene drive system for regional population modification. Nat Commun 2020;11(1):1082; doi: 10.1038/s41467-020-14960-3.

18. Johnson ML, Hay BA, Maselko M. Allele Sails: Launching Traits and Fates into Wild Populations with Mendelian DNA Sequence Modifiers. 2024;2024.03.18.585647; doi: 10.1101/2024.03.18.585647.

19. Teem JL, Alphey L, Descamps S, et al. Genetic Biocontrol for Invasive Species. Frontiers in Bioengineering and Biotechnology 2020;8:452; doi: 10.3389/fbioe.2020.00452.

20. Kandul NP, Liu J, Sanchez C. HM, et al. Transforming insect population control with precision guided sterile males with demonstration in flies. Nat Commun 2019;10(1):84; doi: 10.1038/s41467-018-07964-7.

21. Beach SJ, Maselko M. Recombinant venom proteins in insect seminal fluid reduce female lifespan. Nature Communications 2025;16(1):219; doi: 10.1038/s41467-024-54863-1.

22. Thomas DD, Donnelly CA, Wood RJ, et al. Insect Population Control Using a Dominant, Repressible, Lethal Genetic System. Science 2000;287(5462):2474–2476; doi: 10.1126/science.287.5462.2474.

23. Philips JG, Martin-Avila E, Robold AV. Horizontal gene transfer from genetically modified plants - Regulatory considerations. Front Bioeng Biotechnol 2022;10; doi: 10.3389/fbioe.2022.971402.

24. Thygesen P. Regulation of genome edited organisms in Australia. Transgenic Res 2024;33(6):545–550; doi: 10.1007/s11248-024-00411-y.

25. Shine R. The ecological impact of invasive cane toads (*Bufo marinus*) in Australia. Q Rev Biol 2010;85(3):253–291.

26. Phillips BL, Brown GP, Webb JK, et al. Invasion and the evolution of speed in toads. Nature 2006;439(7078):803–803; doi: 10.1038/439803a.

27. Phillips BL, Brown GP, Shine R. Life-history evolution in range-shifting populations. Ecology 2010;91(6):1617–1627; doi: 10.1890/09-0910.1.

28. Shine R. Invasive species as drivers of evolutionary change: cane toads in tropical Australia. Evol Appl 2012;5(2):107–116; doi: 10.1111/j.1752-4571.2011.00201.x.

29. Lever C. The Cane Toad: The History and Ecology of a Successful Colonist. Westbury Academic and Scientific Publishing: Otley, UK; 2001.

30. Shine R, Ward-Fear G, Brown GP. A famous failure: Why were cane toads an ineffective biocontrol in Australia? Conservation Science and Practice 2020;2(12):e296; doi: 10.1111/csp2.296.

31. Urban MC, Phillips BL, Skelly DK, et al. A toad more traveled: the heterogeneous invasion dynamics of cane toads in Australia. Am Nat 2008;171(3):E134–E148; doi: 10.1086/527494.

32. Jolly CJ, Kelly E, Gillespie GR, et al. Out of the frying pan: Reintroduction of toad-smart northern quolls to southern Kakadu National Park. Austral Ecol 2018;43(2):139–149; doi: 10.1111/aec.12551.

33. Fukuda Y, Tingley R, Crase B, et al. Long-term monitoring reveals declines in an endemic predator following invasion by an exotic prey species. Anim Conserv 2016;19(1):75–87; doi: 10.1111/acv.12218.

34. Phillips BL, Greenlees MJ, Brown GP, et al. Predator behaviour and morphology mediates the impact of an invasive species: cane toads and death adders in Australia. Animal Conservation 2010;13(1):53–59; doi: 10.1111/j.1469-1795.2009.00295.x.

35. Phillips BL, Shine R. An invasive species induces rapid adaptive change in a native predator: cane toads and black snakes in Australia. Proc R Soc B 2006;273(1593):1545–1550; doi: 10.1098/rspb.2006.3479.

36. Pettit L, Somaweera R, Kaiser S, et al. The impact of invasive toads (Bufonidae) on monitor lizards (Varanidae): An overview and prospectus. The Quarterly Review of Biology 2021;96(2):105–125; doi: 10.1086/714483.

37. Price-Rees SJ, Brown GP, Shine R. Predation on toxic cane toads (*Bufo marinus*) may imperil bluetongue lizards (*Tiliqua scincoides intermedia*, Scincidae) in tropical Australia. Wildlife Research 2010;37(2):166; doi: 10.1071/WR09170.

38. Greenlees MJ, Harris S, White AW, et al. The establishment and eradication of an extra-limital population of invasive cane toads. Biol Invasions 2018;20(8):2077–2089; doi: 10.1007/s10530-018-1681-8.

39. Somaweera R, Somaweera N, Shine R. Frogs under friendly fire: How accurately can the general public recognize invasive species? Biological Conservation 2010;143(6):1477– 1484; doi: 10.1016/j.biocon.2010.03.027.

40. Ward-Fear G, Rangers B, Bruny M, et al. Teacher toads: Buffering apex predators from toxic invaders in a remote tropical landscape. Conserv Lett 2024;17(3):e13012; doi: 10.1111/conl.13012.

41. Shine R, Baeckens S. Rapidly evolved traits enable new conservation tools: perspectives from the cane toad invasion of Australia. Evolution 2023;77(8):1744–1755; doi: 10.1093/evolut/qpad102.

42. Crossland MR, Shine R. Cues for cannibalism: cane toad tadpoles use chemical signals to locate and consume conspecific eggs. Oikos 2011;120(3):327–332; doi: 10.1111/j.1600-0706.2010.18911.x.

43. DeVore JL, Crossland MR, Shine R, et al. The evolution of targeted cannibalism and cannibal-induced defenses in invasive populations of cane toads. PNAS 2021;118(35):e2100765118; doi: 10.1073/pnas.2100765118.

44. Doudna JA, Charpentier E. The new frontier of genome engineering with CRISPR-Cas9. Science 2014;346(6213):1258096; doi: 10.1126/science.1258096.

45. Jinek M, Chylinski K, Fonfara I, et al. A Programmable Dual-RNA–Guided DNA Endonuclease in Adaptive Bacterial Immunity. Science 2012;337(6096):816–821; doi: 10.1126/science.1225829.

46. Anders C, Niewoehner O, Duerst A, et al. Structural basis of PAM-dependent target DNA recognition by the Cas9 endonuclease. Nature 2014;513(7519):569–573; doi: 10.1038/nature13579.

47. Chakrabarti AM, Henser-Brownhill T, Monserrat J, et al. Target-Specific Precision of CRISPR-Mediated Genome Editing. Molecular Cell 2019;73(4):699–713.e6; doi: 10.1016/j.molcel.2018.11.031.

48. van Overbeek M, Capurso D, Carter MM, et al. DNA Repair Profiling Reveals Nonrandom Outcomes at Cas9-Mediated Breaks. Molecular Cell 2016;63(4):633–646; doi: 10.1016/j.molcel.2016.06.037.

49. Tuladhar R, Yeu Y, Tyler Piazza J, et al. CRISPR-Cas9-based mutagenesis frequently provokes on-target mRNA misregulation. Nature Communications 2019;10(1):4056; doi: 10.1038/s41467-019-12028-5.

50. Kroll F, Powell GT, Ghosh M, et al. A simple and effective F0 knockout method for rapid screening of behaviour and other complex phenotypes. Ekker SC, Stainier DY, Balciunas D. eds. eLife 2021;10:e59683; doi: 10.7554/eLife.59683.

51. Zuo E, Cai Y-J, Li K, et al. One-step generation of complete gene knockout mice and monkeys by CRISPR/Cas9-mediated gene editing with multiple sgRNAs. Cell Research 2017;27(7):933–945; doi: 10.1038/cr.2017.81.

52. Wang F, Shi Z, Cui Y, et al. Targeted gene disruption in Xenopus laevis using CRISPR/Cas9. Cell & bioscience 2015;5:1–5.

53. Guo X, Zhang T, Hu Z, et al. Efficient RNA/Cas9-mediated genome editing in Xenopus tropicalis. Development 2014;141(3):707–714; doi: 10.1242/dev.099853.

54. Elewa A, Wang H, Talavera-López C, et al. Reading and editing the Pleurodeles waltl genome reveals novel features of tetrapod regeneration. Nature Communications 2017;8(1):2286; doi: 10.1038/s41467-017-01964-9.

55. Flowers GP, Timberlake AT, Mclean KC, et al. Highly efficient targeted mutagenesis in axolotl using Cas9 RNA-guided nuclease. Development 2014;141(10):2165–2171; doi: 10.1242/dev.105072.

56. Sanor LD, Flowers GP, Crews CM. Multiplex CRISPR/Cas screen in regenerating haploid limbs of chimeric Axolotls. Harvey RP, Wittkopp PJ. eds. eLife 2020;9:e48511; doi: 10.7554/eLife.48511.

57. Nakajima K, Nakajima T, Takase M, et al. Generation of albino enopus tropicalis using zinc-finger nucleases. Development, Growth & Differentiation 2012;54(9):777–784; doi: 10.1111/dgd.12006.

58. Nakayama T, Fish MB, Fisher M, et al. Simple and efficient CRISPR/Cas9-mediated targeted mutagenesis in Xenopus tropicalis. Genesis 2013;51(12):835–843; doi: 10.1002/dvg.22720.

59. Gosner KL. A Simplified Table for Staging Anuran Embryos and Larvae with Notes on Identification. Herpetologica 1960;16(3):183–190.

60. Wlizla M, McNamara S, Horb ME. Generation and care of Xenopus laevis and Xenopus tropicalis embryos. Methods in molecular biology (Clifton, NJ) 2018;1865:19; doi: 10.1007/978-1-4939-8784-9_2.

61. Wu RS, Lam II, Clay H, et al. A Rapid Method for Directed Gene Knockout for Screening in G0 Zebrafish. Developmental Cell 2018;46(1):112–125.e4; doi: 10.1016/j.devcel.2018.06.003.

62. Lande R, Arnold SJ. The Measurement of Selection on Correlated Characters. Evolution 1983;37(6):1210–1226; doi: 10.1111/j.1558-5646.1983.tb00236.x.

63. Barrett RDH, Hoekstra HE. Molecular spandrels: tests of adaptation at the genetic level. Nat Rev Genet 2011;12(11):767–780; doi: 10.1038/nrg3015.

64. Cleves PA, Tinoco AI, Bradford J, et al. Reduced thermal tolerance in a coral carrying CRISPR-induced mutations in the gene for a heat-shock transcription factor. Proceedings of the National Academy of Sciences 2020;117(46):28899–28905; doi: 10.1073/pnas.1920779117.

65. Selechnik D, Richardson MF, Shine R, et al. Increased Adaptive Variation Despite Reduced Overall Genetic Diversity in a Rapidly Adapting Invader. Front Genet 2019;10:1221; doi: 10.3389/fgene.2019.01221.

66. Hietpas RT, Jensen JD, Bolon DNA. Experimental illumination of a fitness landscape. Proceedings of the National Academy of Sciences 2011;108(19):7896–7901; doi: 10.1073/pnas.1016024108.

67. Lobkovsky AE, Koonin EV. Replaying the Tape of Life: Quantification of the Predictability of Evolution. Front Genet 2012;3:246; doi: 10.3389/fgene.2012.00246.

